# Identification of methylation and hydroxymethylation haplotype blocks as distal regulatory elements aids in deconvolution of heterogeneous brain tissues

**DOI:** 10.1101/237024

**Authors:** Qin Ma, Zhengzheng Xu, Huan Lu, Ziying Xu, Yuanyuan Zhou, Bifeng Yuan, Weimin Ci

## Abstract

5-Hydroxymethylcytosine (5hmC) is an oxidation product of 5-methylcytosine (5mC), and adjacent CpG sites in the mammalian genome can be co-methylated and co-hydroxymethylated due to the processivity of DNMT and TET enzymes. We applied TAB-seq and oxBS-seq to selectively detect 5hmC and 5mC, respectively, at base resolution in the mouse cortex, olfactory bulb and cerebellum tissues. We found that majority of the called 5hmC CpG sites frequently had 5mC modification simultaneously and enriched in gene body regions of neuron development related genes in brain tissues. These results supported a prominent role of oxidizing 5mC to 5hmC as new epigenetic mark. Strikingly, by a systematic search of regions that show highly coordinated methylation and hydroxymethylation (MHBs and hMHBs), we found that MHBs significantly overlapped with hMHBs in gene body regions which further supported that oxidized 5mC to 5hmC co-ordinately in a subset of cells within heterogeneous brain tissues. Consistently, using a metric called methylation haplotype load, we defined a subset of 1,361 tissue-specific MHBs and 3,818 shared MHBs which were predominantly regulatory elements, and aids in deconvolution of heterogeneous brain tissues. Our results provide new insights into the role of co-ordinately oxidized 5mC to 5hmC as distal regulatory elements may involve in regulating tissue identity.

## Introduction

In contrast with global DNA methylation levels, which are stable across tissues, the levels of 5hmC are highly tissue specific and are abundant in brain tissue (Globisch et al. 2010). The oxidation of 5mC to 5hmC is carried out by the ten–eleven translocation (TET) enzymes (Tahiliani et al. 2009). A number of pathways for the removal of the methyl group from 5mC via 5hmC have been suggested and validated (He et al. 2011; Shen et al. 2013), and thus, 5hmC is proposed as an intermediate of DNA demethylation. However, given that 5hmC binds to specific regulatory proteins (Mellen et al. 2012; Spruijt et al. 2013) and that it is mainly present at actively transcribed genes (Kriaucionis and Heintz 2009; Szulwach et al. 2011; Mellen et al. 2012), it appears that 5hmC may convey regulatory functions and be an epigenetic mark on its own. Additionally, a previous study showed that regions with low methylation (LMRs) demonstrate features of distal regulatory elements and correspond to cell type-specific transcription factor binding sites (Stadler et al. 2011). Additionally, the oxidization of 5mC to 5hmC occurs preferentially at LMRs (Feldmann et al. 2013). These results indicated that the tissue-specific oxidization of 5mC to 5hmC might facilitate the establishment of tissue-specific regulatory elements.

The identification of the tissue-specific 5hmC pattern can be harnessed for analysing the relative cell composition of heterogeneous samples, such as brain tissues, because they exhibit cell-type-specific 5hmC patterns. Due to the locally coordinated activities of DNMT and TET dioxygenase proteins, adjacent CpG sites on the same DNA molecule can share similar methylation and hydroxymethylation statuses. Thus, the analysis of CpG co-methylation and co-hydroxymethylation in cell populations within a tissue may aid in the deconvolution of heterogeneous tissue samples and elucidate the key regulatory elements for tissue identity. Therefore, the theoretical framework of linkage disequilibrium (Slatkin 2008), which was developed to model the co-segregation of adjacent genetic variants on chromosomes in human populations, can be applied to the analysis of CpG co-methylation and co-hydroxymethylation in the cell populations of a tissue. Consistently, a recent study applied base-resolution whole-genome bisulfite sequencing (BS-seq) data across the largest set of human tissue types with a metric called methylation haplotype load (MHL) in highly coordinated methylation blocks (MHBs). The study showed that a subset informative blocks aids in the deconvolution of heterogeneous tissue samples and predicts the tissue-of-origin from plasma DNA (Guo et al. 2017). These results support that the same theoretical framework can be applied to model both the co-methylation and co-hydroxymethylation of adjacent CpG sites. However, in the only computationally inferred from a combination of different datasets, such as BS-seq (5mC + 5hmC) and TAB-seq (5hmC). However, the co-occurrence of 5mC and 5hmC on the same cytosine within a population of cells, and the co-methylation and co-hydroxymethylation of adjacent CpG sites cannot be determined from these datasets. Single-base-resolution profiling of both 5mC and 5hmC is fundamental to understanding the function role of the turnover from 5mC to 5hmC or stable methylation.

To the best of our knowledge, TAB-seq (Tet-assisted bisulfite sequencing) (Yu et al. 2012) or oxBS-seq (oxidative bisulfite sequencing) (Booth et al. 2013) are the only currently available methods that can selectively detect 5hmC and 5mC, respectively, at base resolution. Herein, we applied the oxBS-seq and TAB-seq to profile 5mC and 5hmC at the single-nucleotide level in three brain tissues, including the cortex, olfactory bulb and cerebellum. Our results supported that a prominent role of oxidizing 5mC to 5hmC as new epigenetic marks, and co-ordinately oxidized 5mC to 5hmC as distal regulatory elements may involve in regulating tissue identity.

## Results

### Oxidization 5mC to 5hmC, generating new epigenetic marks in brain tissues

Firstly, we performed TAB-seq and oxBS-seq on cerebellum, cortex and olfactory bulb tissues derived from female mice. Biological replicates were included for each tissue sample from an independent mouse. The hierarchical clustering clearly showed that both hydroxymethylomes and methylomes stored memory of the tissue identities with greatly concordant among replicates (Supplemental Figure S1A). We sequenced to an average depth of 23-fold genomic coverage per tissue (Supplemental Table S1). Consistent with previous study, we observed non-CpG methylation in all three brain tissues but the majority of 5hmC and 5mC exists in a CpG context (Supplemental Figure S1B and S1C). Next, we focused on the 5hmC and 5mC modifications in the CpG context. As exampled in the *Auts2* gene locus, although the overall modification (5mC+5hmC) was similar among the three brain tissues as determined by the BS-seq method(Hon et al. 2013), the extent and/or distribution of oxidized 5mC to 5hmC were different among tissues (Figure 1A). The relative 5mC and 5hmC levels in the three tissues are consistent with the results from mass spectrometry (Supplemental Figure S1D).

**Figure 1.**
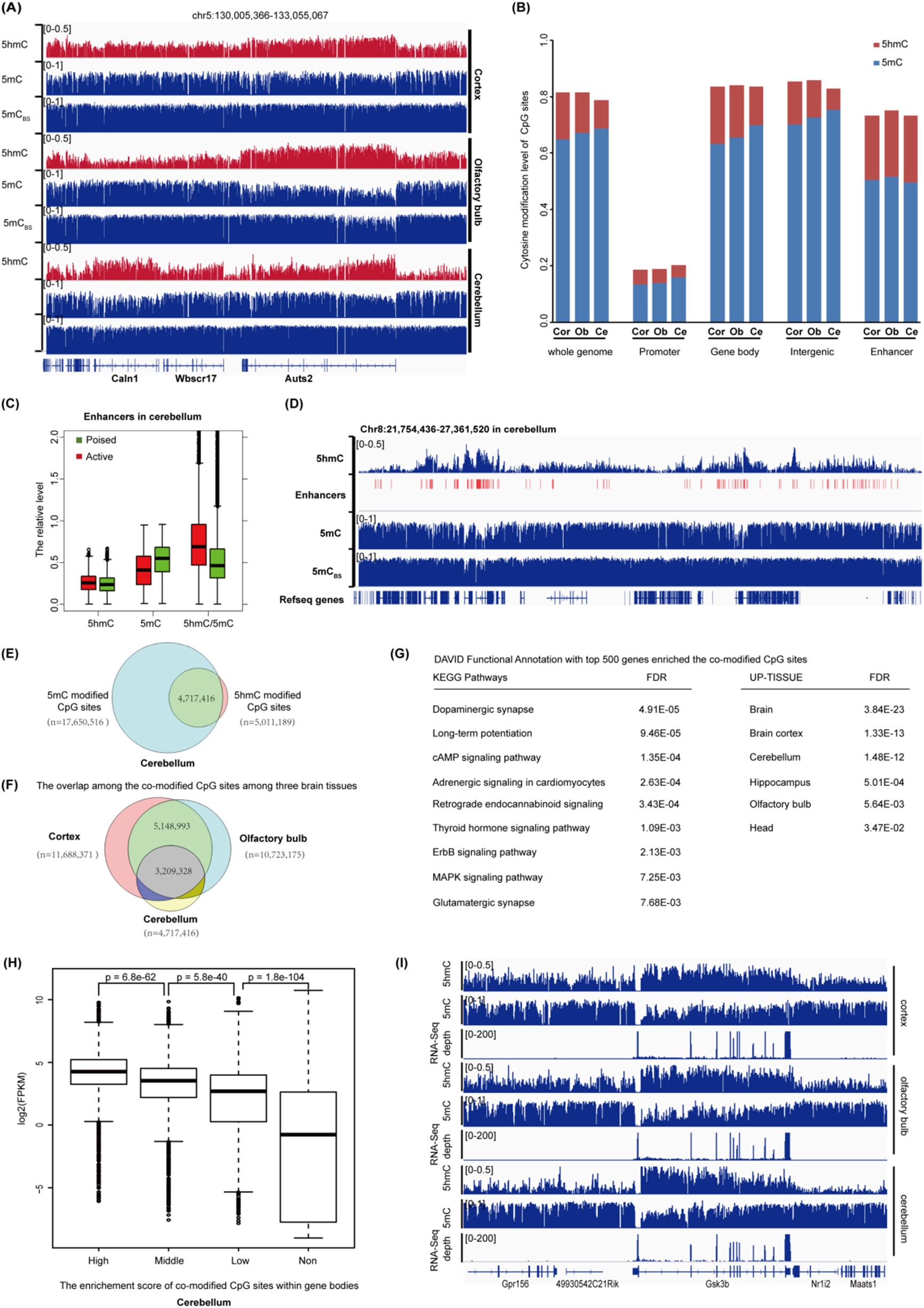
Oxidization 5mC to 5hmC occurs globally at gene bodies for brain-development-related genes. **(A)** Wiggle tracks of the 5hmC, 5mC, and total modification (5hmC + 5mC) profiles in a genomic region on chromosome 5. **(B)** Average 5hmC and 5mC levels in different genomic elements determined by TAB-seq and oxBS-seq, respectively. The promoter is defined as being 500 bp upstream of the TSS. **(C)** Box plots of the 5hmC, 5mC and 5hmC/5mC levels in the active enhancers and poised enhancers in the cerebellum. **(D)** Wiggle tracks of the 5hmC, 5mC, and total modification (5hmC + 5mC) profiles for a representative region with enriched enhancers. **(E)** Venn plot of the co-occurrence of 5hmC and 5mC at the same CpG site in the cerebellum. **(F)** Venn plot of the overlap for the co-occurrence CpG sites among the three brain tissues. **(G)** The DAVID Functional Annotation for the top 500 genes enriched in the overlapped CpG sites among the three brain tissues. **(H)** The association between gene expression level and the enrichment score of the co-modified CpG sites within gene body regions. The enrichment score was calculated as described in ‘Materials and Methods’ sections. Non represents that the enrichment score is less than 0. Statistical significances were evaluated by rank-sum test. **(I)** Graphical representation of the association between gene expression (depths of RNA-seq) and the average 5hmC, 5mC level in *gsk3b* gene locus.

Moreover, the co-occurrence of 5mC and 5hmC is identified in all genomic elements, especially enhancers (Figure 1B). We found that, in contrast to the relative depletion of 5mC, 5hmC CpG sites were enriched in the enhancer regions (Figure 1B, Supplemental Figure S1E), suggesting that the oxidation of 5mC to 5hmC may play a role in regulating enhancer activity. Previous study with BS-seq method has shown that tissue-specific differentially methylated regions are hypomethylated and predominantly localized to distal cis-regulatory elements especially enhancers. Unexpectedly, some of these ‘vestigial’ enhancers lack DNA methylation in adult tissue but remain inactive (Hon et al. 2013). Thus, oxidization of 5mC to 5hmC but not DNA demethylation (hypomethylation) may also be required for enhancer activity which will escape from the detection of BS-seq method (Huang et al. 2010; Jin et al. 2010). To address this, we compiled the available enhancer data for the three brain tissues from the mouse Encyclopedia of DNA Elements (ENCODE) Project (Shivakumar et al. 2002). As predicted, although the global hypomethylation can significantly distinguish the active enhancers from poised enhancers, active enhancers also showed significantly higher levels of 5hmC/5mC compared to poised enhancers (Figure 1C). A representative locus is shown in Figure 1D. Thus, oxidized 5mC to 5hmC may play an important role in tissue identity, at least partially through the regulation of enhancer activity.

More importantly, with the single-nucleotide resolution mappings of both 5mC and 5hmC, we found that majority of the called 5hmC CpG sites frequently have 5mC modification simultaneously (co-modified sites) in all three brain tissues (Figure 1E, Supplemental Figure S1F). Given that 5mC and 5hmC are binary for any particular cytosine, the co-occurrence of the two modifications indicates that the oxidization of 5mC to 5hmC occurs in a subset of cells within the tissues. Thus, oxidization 5mC to 5hmC may generate new epigenetic marks in regulating brain tissue identity. Consistent with this scenario, we found that the co-modified CpG sites were not randomly distributed in brain tissues and showed great concordance among three brain tissues (Figure 1F). Furthermore, DAVID functional annotation for the genes enriched the shared co-modified sites among the three tissues revealed that multiple neuron development related KEGG pathways, such as dopaminergic synapse, retrograde endocannabinoid signaling and Glutamatergic synapse (Figure 1G). And these genes were also specifically expressed in the brain (Figure 1G, DAVID Functional Annotation, UP-TISSUE). More importantly, further analyses including RNA-seq data showed that the genes enriched co-modified CpG sites within gene body regions expressed higher compared to the genes without enrichments (Figure 1H, Supplemental Figure S1G). A representative locus at *GSK3B* gene, a serine-threonine kinase involved in neuronal cell development (Kwok et al. 2008), was shown in Figure 1i. Collectively, oxidization of 5mC to 5hmC generated new epigenetic marks which may regulate the brain tissue identity.

### Identification of methylation and hydroxymethylation haplotype blocks that significantly overlapped and co-localized with the regulatory elements

Since both 5mC and 5hmC are cell-type specific, and the functional regions can be harnessed for analysing the relative cell composition of heterogeneous tissues. Most of the recent efforts rely on the 5mC and 5hmC level of individual CpG sites, and they are limited by the technical noise and sensitivity in measuring single CpG modification level especially for the co-modification sites. However, if the oxidization is coordinated, the adjacent CpG sites should be co-methylated/co-hydroxymethylated due to the processivity of DNMTs and TETs. Thus, we applied the same concept of genetic linkage disequilibrium (Slatkin 2008) and the *r^2^* metric. An unbiased methylation status of multiple CpG sites in paired Illumina sequencing reads of oxBS-seq were extracted to form methylation haplotypes, and a pairwise “linkage disequilibrium” of CpG methylation *r^2^* was calculated as previously reported (Guo et al. 2017). We partitioned the genome into blocks of tightly coupled CpG methylation sites (which we refer to as MHBs; a representative MHB is shown in Figure 2A) using an *r^2^* cutoff of 0.5. We identified 10,809 MHBs with an average size of 501 bp and a minimum of four CpGs per block, which tended to be tightly co-regulated on the epigenetic status at the level of single DNA molecules (Supplemental Figure S2A, B; Supplemental Table S2). The majority of the CpG sites within the same MHBs were nearly perfectly coupled (*r^2^* ~ 1.0). The MHBs established by the oxBS-seq data represent a distinct type of genomic feature that partially overlaps with known genomic elements. Among all of the MHBs, 4,421 (40.9%) were located in intergenic regions, whereas 6,388 (59.1%) regions were located in transcribed regions (Figure 2B). These MHBs were significantly enriched in promoters, gene body regions, and enhancers (Figure 2C). Therefore, MHBs probably capture the local coherent epigenetic signatures that are directly or indirectly coupled to the regulatory elements. To further prove the oxidization of 5mC to 5hmC occurring in coordinated manner, we examined whether the MHBs overlapped with hydroxymethylation haplotype blocks (hMHBs) defined by the TAB-seq data (see ‘Materials and Methods’ section). As expected, we did find that the hMHBs were significantly overlapped with MHBs in gene body regions (Figure 2D, E). Collectively, co-ordinately oxidation 5mC to 5hmC at block level may generate the regulatory elements for tissue identity.

**Figure 2.**
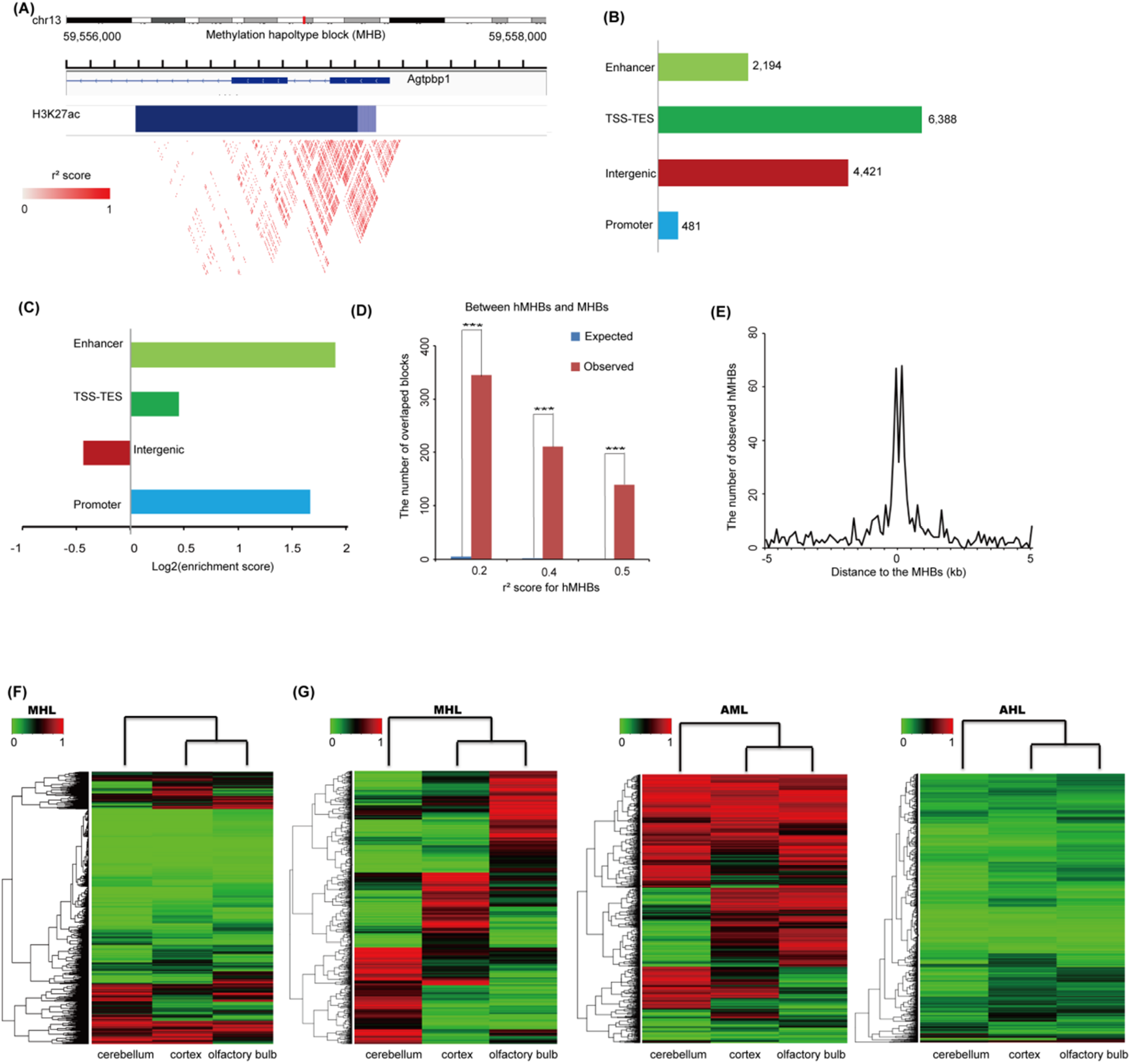
Characterization of the methylation haplotype blocks (MHBs) and tissue clustering based on methylation haplotype load. **(A)** An example of a MHB at the gene body of the gene *Agtpbp1*. The r^2^ scores between the CpG sites in the MHB were represented in heatmap mode using the WashU EpiGenome Browser. The scale bar from white to red represents the r^2^ scores between the two CpG sites from low to high. **(B)** Co-localization of MHBs (n=10,809) with known genomic elements. **(C)** Enrichment of MHBs in known genomic features. **(D)** The coordination between the hydroxymethylation haplotype blocks (hMHBs) and MHBs in gene body regions. The expected number of overlapped blocks was evaluated by random sampling method. The coordination was measured under different cut-offs for the *r^2^* score of linkage disequilibrium two adjacent CpG sites hydroxymethylation. *P* values were calculated by hyper-geometric test. *** represents *p* value < 0.0001. **(E)** The observed number of the segment middle for hMHBs within ±5k of the segment middle for MHBs within gene body regions. The number for the segment middle for hMHBs was evaluated in the 100 bp bin. **(F)** MHL-based unsupervised clustering of mouse cerebellum, cortex and olfactory bulb using all MHBs. **(G)** Comparison of the cluster performance to the brain tissues using the MHL, the average methylation level (AML) and the average hydroxymethylation level (AHL) metrics in the tissue-specific MHBs. The MHL exhibits a better signal-to-noise ratio than the AML and AHL for tissue clustering.

### Block-level analysis using methylation haplotype load can aid in the deconvolution of heterogeneous brain tissues

To enable a quantitative analysis of the co-methylation patterns within individual MHBs across samples, we used the same metric called methylation haplotype load (MHL) as the weighted mean of the fraction of fully methylated haplotypes and substrings at different lengths (Guo et al. 2017). It has been shown that MHL is capable of distinguishing blocks that have the same average levels of methylation but various degrees of coordinated methylation (Guo et al. 2017). In addition, the MHL is bounded between 0 and 1, which allows for a direct comparison of different regions across many datasets.

Consistent with a previous study (Guo et al. 2017), the block-level analysis based on the MHL distinguished these three brain tissues, although the three brain tissue types shared rather similar cell compositions (Figure 2F). To identify a subset of MHBs for effective clustering of these three brain tissues, we calculated a tissue specific index (TSI) for each MHB (Materials and Methods). We identified a set of 1,361 tissue-specific MHBs and 3,818 shared MHBs (Supplemental Table S3). Using these tissue-specific MHBs, we compared the performance for effective clustering of the different tissues between the MHL, the average methylation level (AML) and the average hydroxymethylation level (AHL) in the MHBs. The MHL provided the best tissue identity, which might be due to the higher biological or technical variability of the sequencing methods for 5mC and 5hmC, such as incomplete bisulfide conversion, TET oxidation rate or sequencing errors (Figure 2G).

### Tissue-specific MHBs but not shared MHBs capture the local co-modified regions

Since oxidization of 5mC to 5hmC has been suggested to be involved in tissue identity (Stadler et al. 2011; Feldmann et al. 2013), we next explored whether the block-level analysis based on the MHL captures the local co-modified regions, especially in the tissue-specific MHBs. Strikingly, we found that the tissue-specific MHBs (n=1,361) have a high MHL, the highest level of 5hmC and moderate levels of 5mC, which were specifically enriched in the enhancer regions (Figure 3A, D; Supplemental Figure S3A, B). This is consistent with a previous finding showing that regions with low methylation (LMRs) demonstrate features of distal regulatory elements (Stadler et al. 2011), and the oxidization of 5mC to 5hmC occurs preferentially at LMRs (Feldmann et al. 2013). In contrast, the shared MHB showed a bimodal distribution of the MHL (1,257 for MHB with high MHL, 2,561 for MHB with low MHL). The shared MHBs with low MHLs had both the lowest 5mC and 5hmC levels, which were enriched in promoter regions (Figure 3B, D). In contrast, the shared MHBs with high MHLs had the highest 5mC level and a residual 5hmC level, which were enriched in the intergenic regions (Figure 3C, D). Consistently, we identified a significant enrichment of co-modified CpG sites in the tissue-specific MHBs but not in the shared MHBs in all three tissues (Figure 3E). Collectively, tissue-specific MHBs capture the local co-modified regions, which may represent tissue-specific regulatory elements.

**Figure 3.**
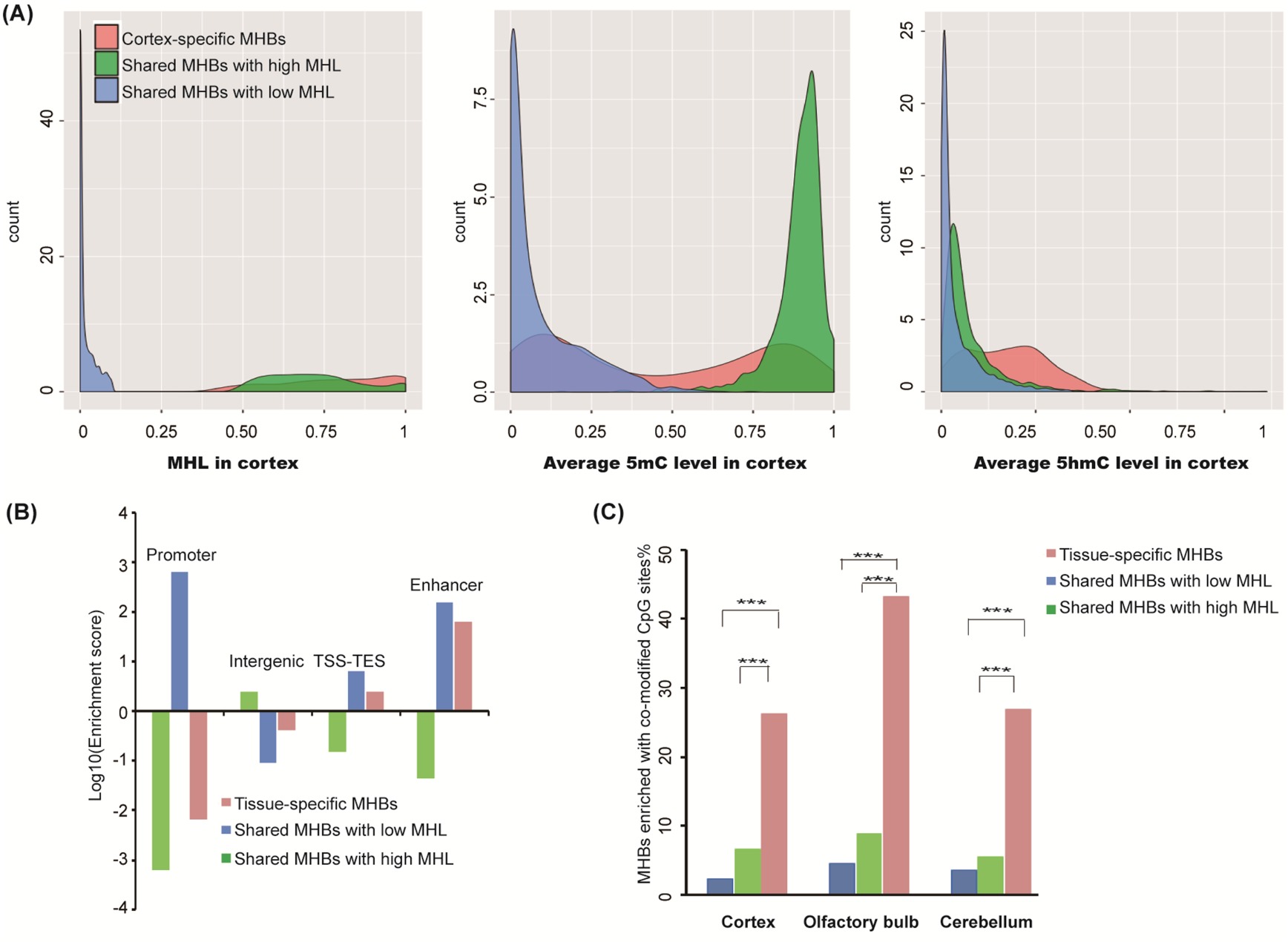
Tissue-specific co-methylation regions capture the local co-hydroxymethylation regions. **(A)** The density plots of the MHL and 5mC and 5hmC levels in tissue-specific MHBs and the shared MHBs with high MHLs or low MHLs in the cortex tissue. **(B)** Heatmap representing the MHL and 5mC and 5hmC levels in the shared MHBs with low MHLs. **(C)** Heatmap representing the MHL and 5mC and 5hmC levels in the shared MHBs with high MHLs. **(D)** Enrichment of tissue-specific MHBs and shared MHBs with low or high MHLs in known genomic features. **(E)** The percentages of MHBs with enriched co-modified CpG sites for tissue-specific MHBs and shared MHBs with low or high MHLs in the three tissues. Statistical significance is evaluated by hyper-geometric test, *** represents *p* value < 0.0001.

### Shared MHBs with low MHL, but not tissue-specific MHBs, correspond with developmental enhancers

Previous study of the methylomes across 17 mice tissues including cortex, olfactory bulb and cerebellum with BS-seq method has generated a large number of tissue-specific differentially methylated regions (tsDMRs) that are hypomethylated in a tissue-specific manner (Hon et al. 2013). A significant percentage of these tsDMRs correspond with dormant developmental enhancers (2646 of 7761 in cortex, 6667 of 16086 in olfactory bulb and 15410 of 29558 in cerebellum). Then, we compared them to publicly available genomic annotations. We found that shared MHBs with low MHL were enriched in active enhancers in embryonic stem cells and developing brain from E14.5, and shared MHBs with high MHL were not enriched in active enhancers (Figure 4A and 4B). These results are consistent with recent findings that some hypomethylated regulatory elements are dormant in adult tissue but active in embryonic development (Hon et al. 2013). In contrast, tissue-specific MHBs were enriched for stronger enhancers in adult tissues compared to embryonic stem cells and the whole brain from embryonic day (E) 14.5. Considering the higher level of active chromatin modifications is identified in shared MHBs with low MHL compared to tissue-specific MHBs (Figure 4A and 4B), we further explored the possibility that the tissue-specific MHBs identified here may represent a new set of tissue-specific regulatory elements which may escape from the current detection method, such as BS-seq. Collectively, shared MHBs with low MHL, but not tissue-specific MHBs, correspond with developmental enhancers.

**Figure 4.**
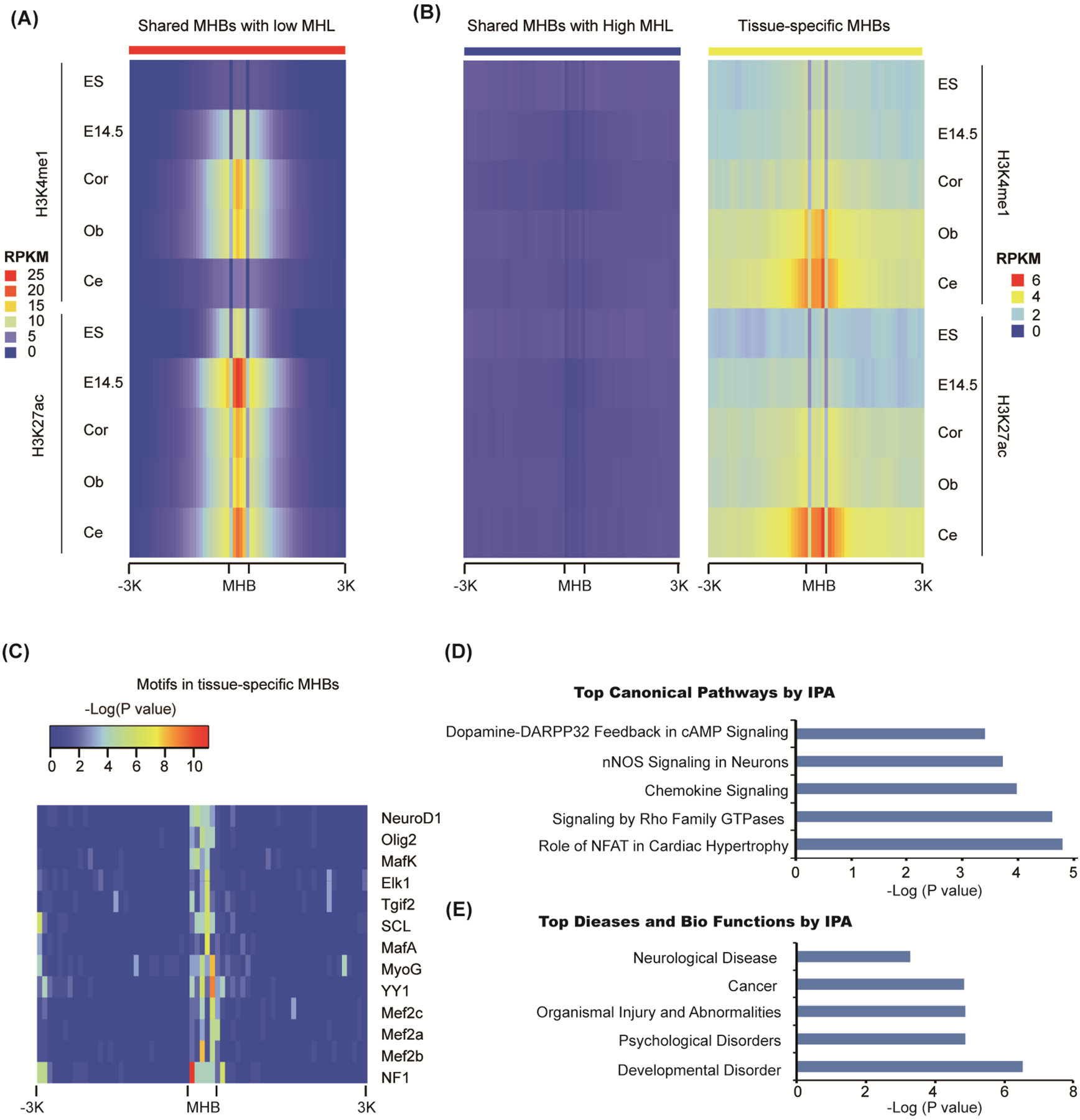
Tissue-specific MHBs resembled the regulatory elements for tissue identity. **(A)** Heatmap representing the H3K4me1 and H3K27ac enrichment (reads per kilobase per million mapped reads (RPKM)) around the shared MHBs with low MHL in adult tissues, mouse embryonic stem cells, and the whole brain tissue from E14.5 embryo. Cor, Ob and Ce represent cortex, olfactory bulb and cerebellum tissues respectively. **(B)** The enrichment of H3K4me1 and H3K27ac around the shared MHBs with high MHL and tissue-specific MHBs. **(C)** Transcriptional factor binding motif enrichments near tissue-specific MHBs. **(D)** and **(E)** The IPA analyses of the genes overlapped with tissue-specific MHBs found canonical pathways and diseases significantly affected.

### Tissue-specific MHBs resembled the regulatory elements for tissue identity

Next, we investigated the possibility that the MHBs identified here might correspond with distal cis-regulatory elements themselves. Thus, we speculated that tissue-specific MHBs might capture binding events of transcription factors specific to tissue identity compared to the shared MHBs. Indeed, consensus motifs for known brain-specific master regulators were significantly enriched in the tissue-specific MHBs. For example, motifs for neuronal development factors NeuroD1, Olig2 and MEF2A, B, and D were specifically identified (Figure 4C). All of these factors are involved in synaptic differentiation and neural crest differentiation (de Melo et al. 2011; Zembrzycki et al. 2015).

Next, we used the Ingenuity Pathway Analysis (IPA) to predict functions of the genes overlapped with tissue-specific MHBs. As expected, multiple neuron development related pathways were identified, such as Dopamine-DARPP32 Feedback in cAMP Signaling and nNOS Signaling in Neurons (Figure 4D). And further top diseases and bio functions also supported these genes may involve in neurological disease and cancer (Figure 4E).

More importantly, we also used the GREAT tool (McLean et al. 2010) to predict functions of cis-regulatory region that is the cortex-specific, olfactory bulb-specific and cerebellum-specific MHBs. With limited regions (396 for cortex-specific, 508 for olfactory bulb-specific and 457 for cerebellum-specific), the GO Biological Processes and Cellular Components related with the tissue identity were identified. For example, the GO Biological Processes enriched in the cortex-specific MHBs included the regulation of synaptic transmission and the regulation of transmission of nerve impulse (Supplemental Figure S4A). All these biological processes are needed for cortex identity. Moreover, the GO Biological Processes enriched in the cerebellum-specific MHBs included homotypic cell-cell adhesion, which plays a crucial role in migration, the pathfinding of axons, and synaptic plasticity (Togashi et al. 2009) (Figure 4D). Additionally, the GO cellular component analysis showed that the PML body and microtubule cytoskeleton in cell cortex part (the region of a cell that lies just beneath the plasma membrane) were enriched in the cortex-specific MHBs and the olfactory bulb-specific MHBs, respectively (Supplemental Figure S4B). These results are consistent with previous findings that PML is broadly expressed across the grey matter, with the highest levels in the cerebral and cerebellar cortices (Hall et al. 2016). Additionally, it has been shown that the cytoskeletal organization of the developing mouse olfactory system in the axon is fundamental for olfactory processing (Akins and Greer 2006). Collectively, tissue-specific MHBs corresponded to the regulatory elements for tissue identity.

## Discussion

Herein, with simultaneous single-nucleotide resolution mapping of 5mC and 5hmC, we present the first evidence, at base resolution, that oxidization of 5mC to 5hmC generated new epigenetic markers for a subset of cells within heterogeneous brain tissues. Additionally, since both 5mC and 5hmC are cell-type specific, the identification of tissue-specific differentially methylated or hydroxymethylated regions is harnessed by the complex cell components within the tissues. A recent study showed that the identification of methylation haplotype blocks aids in the deconvolution of heterogeneous tissue (Guo et al. 2017). Herein, we extended the established analysis of co-methylated CpG patterns with the oxBS-seq data. Although the mathematical representations are identical, one major difference is that the oxBS-seq method can unbiasedly measure the 5mC level instead of the sum of 5mC and 5hmC for the BS-seq data. Additionally, we also extended the analysis to TAB-seq data, one major difference is that co-hydroxymethylation analysis must focus on gene body regions. Since the readout of hypermethylated (such as highly methylated intergenic regions) and unmethylated sites (such as promoter regions and CpG islands) will be “T” in TAB-seq data. These regions will false positively called as co-hydroxymethylation regions if included. More importantly, applying the theoretical framework with ox-BS-seq and TAB-seq data, the identified MHBs significantly overlapped with hMHBs, and resembled the regulatory elements for tissue identity. In comparison, neither AML nor AHL within these MHBs can distinguish the tissue identity, which might be due to the higher biological or technical variability of the sequencing methods for 5mC and 5hmC. Thus, these blocks will not be identified with either BS-seq data or the theoretical framework as differentially methylated/hydroxymethylated regions. Consistently, previous study with BS-seq method has shown that tissue-specific differentially methylated regions are hypomethylated and predominantly localized to distal cis-regulatory elements especially enhancers. Unexpectedly, some of these ‘vestigial’ enhancers lack DNA methylation in adult tissue but remain inactive (Hon et al. 2013). Collectively, our results supported that the oxidization of 5mC to 5hmC, without changing the overall modification level of cytosine, plays a pivotal role in regulating tissue identity, which may escape the detection of the BS-seq method.

Furthermore, recent studies (Stadler et al. 2011) propose that epigenetic changes could result from prior events dictated by genetic information, such as TF binding. The methylated DNA binding factors, the potential pioneer factors, may bind chromatin (highly methylated) and turn it into a regulatory element through the turnover to 5hmC (Feldmann et al. 2013). Consistently, a recent study showed that many members of TFs, such as NeuroD1 and the extended homeodomain family, preferred to bind to mCpG-containing sequences. And TFs that preferred mCpG are commonly involved in embryonic and organismal developmental processes (Yin et al. 2017). Strikingly, in our study, NeuroD1 and homeodomain TF, TGIF2, are also identified in the tissue-specific MHBs. Thus, these transcription factors may act as the pioneer factors in establishing the regulatory elements for tissue identity. However, at this point, we do not show that this phenomenon can go beyond correlation. The genetic deletion of the potential pioneer factors would be needed to further support this hypothesis. Investigation is also warranted to determine if the presence of hydroxymethylation is actually involved in the activity of these potential regulatory elements through specific readers.

In summary, by integrating genome-wide single nucleotide resolution sequencings for both 5mC and 5hmC with the theoretical framework of linkage disequilibrium, we have generated a catalogue of distal regulatory elements and tissue-specific co-methylation regions, which capture the local co-modified regions that correspond to the regulatory elements for tissue identity. Additionally, pronounced reciprocal 5mC and 5hmC changes are also identified at cancer-related genes during tumorigenesis (Chen et al. 2016). Thus, the theoretical framework can be applied to identify the biomarkers for cancer-specific pathologies that may be valuable in further understanding cancer biology, diagnosis and therapy. Moreover, it will be vital in future studies to use methods that distinguish 5mC from 5hmC, such as oxBS-seq, when profiling DNA methylation in the genome.

## Materials and methods

### Tissue samples and genomic DNA extraction

The cerebellum, cortex, and olfactory bulb were dissected from an 8-week-old female C57Bl/6 mouse. The cortex tissue came from the whole cerebral cortex excluding the hippocampus regions. The cerebellum and olfactory bulb tissues came from the whole tissues, respectively. The tissues for the technical replicates were from independent mice, and at least two replicates were included in the different assays as indicated. Genomic DNA was isolated using a DNA extraction kit (Qiagen, Cat#: 51306) according to the manufacturers’ instructions.

### LC-ESI-MS analysis

The LC-ESI-MS analysis was performed according to our previous study (Chen et al. 2016). Briefly, the percentages of 5mC and 5hmC were calculated by the following formula, where M(cytosine), M(5mC) and M(5hmC) are the molar quantities of cytosine.

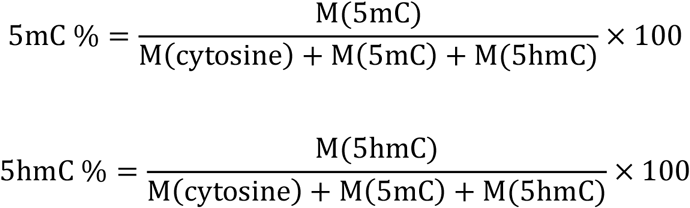

### oxBS-seq

The sequencing libraries were constructed as described (Booth et al. 2013) with minor modifications. Briefly, genomic DNA (0.5 to 1 μg) was end-repaired, A-tailed, and ligated to methylated adaptors following the manufacturer’s instructions. The ligated fragments were then purified using the Bio-Rad Micro Bio-Spin P-6 SSC column (SSC buffer). After the purification, the samples were denatured using 1 M NaOH and oxidized using an oxidant solution (15 mM KRuO_4_ in 0.05 M NaOH). After the purification, the DNA underwent bisulfite conversion using the EZ DNA Methylation Gold kit (Zymo Research) according to the instruction manual. The library was sequenced using Illumina HiSeq X Ten. The paired reads were uniquely mapped to the reference genome (mm10, UCSC) by Bismark. The efficient conversion of 5hmC to uracil was calculated by a spiked 5hmC control from Zymo Research (Cat#: D5405).

### TAB-seq

The sequencing libraries were constructed as described (Yu et al. 2012) with minor modifications. Briefly, genomic DNA, with spike-in controls, was glycosylated and oxidized using the kit from Wisegene (Cat#: K001). Then, the DNA underwent bisulfite conversion using the EZ DNA Methylation Gold kit (Zymo Research) according to the instruction manual. The library was sequenced using Illumina HiSeq X Ten. The paired reads were mapped uniquely to the reference genome (mm10, UCSC) by Bismark. The efficient conversion of unmodified cytosine to uracil and the efficient conversion of 5mC to 5caU/U were calculated using spiked M.SssI-treated lambda DNA.

### The associations between the biological replicates for TAB-seq and oxBS-seq

The 5hmC and 5mC levels of the biological replicates were calculated in 1 Mb regions throughout the genome. Then, a hierarchical cluster analysis was performed using hcluster in R with ward.D methods.

### Quantification of the 5hmC and 5mC levels of each CpG site

The 5hmC level or the 5mC level of each CpG site was calculated using the same methods as described (Chen et al. 2016). Briefly, a binomial distribution model was performed to calculate the significance for the hydroxymethylation or methylation. Only sites with Benjamini-Hochberg corrected binomial *P* value ⩽ 0.05 and reads coverage ⩾ 5 were considered hydroxymethylated or methylated. We counted the number of “C” bases from the sequencing reads as hydroxymethylated or methylated (denoted as N_C_) and the number of “T” bases as unmodified (denoted as N_T_). The hydroxymethylation level or methylation level was estimated as N_C_/(N_C_ + N_T_).

### The enrichment scores of the 5hmC- and 5mC-modified sites in the different genomic regions

The enrichment score was calculated by the following formula: The enrichment score _in the genomic element_ = log2 (# called modified sites _in the genomic element_/# expected). # expected was computed as: # called modified sites _in the genome_ × # CpG sites _in the genomic element_/# total CpG sites _in the genome_. # denotes the number of sites.

### Identification of the genes that enriched the co-modified CpG sites

First, the CpG sites that were simultaneously identified as 5hmC and 5mC modified sites were determined as the co-modified CpG sites. Then, the enrichment score of the sites in each gene was calculated by the following formula:

The enrichment score _in the gene_ = log2 (#the co-modified CpG sites)_in the gene_/# expected). # expected was computed as: # the co-modified CpG sites _in the genome_ × # CpG sites _in the gene_/# total CpG sites _in the genome_. # denotes the number of sites.

### Identification of the methylation haplotype blocks (MHBs)

The MHBs were identified as described (Guo et al. 2017) with minor modifications. Briefly, the mouse genome was split into non-overlapping ‘sequenceable and mappable’ segments using the oxBS-seq data from the mouse cerebellum, cortex and olfactory bulb tissues. The mapped reads from the oxBS-seq datasets were converted into methylation haplotypes within each segment. The methylation linkage disequilibrium was calculated on the combined methylation haplotypes. We then partitioned each segment into MHBs. Candidate MHBs were defined as the genomic region in which the r^2^ value of two adjacent CpG sites was no less than 0.5. Then, the LD r^2^ matrix of the segments was calculated. The candidate MHBs were extended if r_2_ the r^2^ value of the CpG sites in the segments was no less than 0.5.

### Identification of the hydroxymethylation haplotype blocks (hMHBs)

We identified hMHBs using the similar theoretical framework as MHBs with minor modifications. Briefly, for TAB-seq data, the readout of both methylated and unmodified sites will be “T”, while the readout of hydroxymethylated sites will be “C”. Thus, we focused on the gene body regions which enriched 5hmC modifications to identify the co-hydroxymethylation blocks (hMHBs). In this way, we can avoid the false positively called as co-hydroxymethylation blocks for the highly methylated regions (such as intergenic regions) and unmethylated regions (such as promoter regions and CpG islands) with TAB-seq data. Additionally, the average 5hmC level of each called CpG sites is lower than the average 5mC level, thus we tried different cut-offs for the r^2^ score of linkage disequilibrium two adjacent CpG sites hydroxymethylation.

### The coordination between the hMHBs and MHBs in gene body regions

The coordination analysis was performed by random sampling method. First, the same amount of regions with same length distribution as MHBs within gene body regions were randomly sampled and repeated 10,000 times. Then, the overlap between randomly sampled regions and the hMHBs was calculated as the expected number of the overlapped blocks between hMHBs and MHBs within gene body regions. Statistical significance was calculated by hyper-geometric test

### Enrichment analysis of the methylation haplotype blocks for known genomic regulatory elements

The enrichment analysis was performed by random sampling as previously described (Timmons et al. 2015). Genomic regions with the same number of MHBs and CpG numbers were randomly sampled within the mappable CpG regions and repeated 10,000 times. Then, we obtained the expected MHBs in the genomic element. All of the genomic coordinates were based on the mm10 mouse genomic sequence. The enrichment score was calculated by the following formula: The enrichment score _in the genomic element_ = log2 (# called MHBs _in the genomic element_/# expected).

### Methylation haplotype load (MHL)

The MHLs were identified as described (Guo et al. 2017), which is the normalized fraction of methylated haplotypes at different lengths.

### Identification of tissue-specific MHBs

The tissue-specific MHBs were identified as described (Guo et al. 2017) with minor modifications. To investigate the tissue-specific MHBs, the tissue-specific index (TSI) was defined. An empirical threshold TSI > 0.6 was used to define the tissue-specific MHBs.

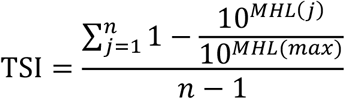

where *n* indicates the number of the tissues, MHL(j) denotes the MHL of *j*^th^ tissue, and MHL max denotes the MHL of the highest methylated tissue.

### Enrichment analysis of the MHBs for the co-modified CpG sites

First, the co-modified CpG sites in each MHB were calculated. Then, the enrichment score of the co-modified CpG sites in each MHB was calculated by the following formula:

The enrichment score _in the MHB_ = log2 (#the co-modified CpG sites)_in the MHB_/# expected). # expected was computed as: # the co-modified CpG sites _in the genome_ × # CpG sites in _the MHB_/# total CpG sites _in the genome_. # denotes the number of sites.

### Motif analysis

We searched for the enrichment of known motifs using the Homer tool (Heinz et al. 2010). To search for the motifs within a single tissue, we used default parameters.

### External datasets

According to the H3K4me1 and H3K27ac marks, two types of enhancers were distinguished as follows: active enhancers that were simultaneously marked by a distal H3K4me1 and H3K27ac and poised enhancers that were solely marked by a distal H3K4me1. The H3K4me1 and H3K27ac peaks of the adult mouse cortex, olfactory bulb and cerebellum tissues were acquired from the ENCODE project.

### Data access

The raw sequence data reported in this paper have been deposited in the Genome Sequence Archive in BIG Data Center (Members 2017), Beijing Institute of Genomics (BIG), Chinese Academy of Sciences, under accession numbers PRJCA000310. The raw data is also deposited in the GEO (GSE97568). The following link was created to allow review of record GSE97568 while it remains in private status: https://www.ncbi.nlm.nih.gov/geo/query/acc.cgi?token=kjwfyiksvfcvvgb&acc=GSE97568.

## Author contributions

The projects were conceived of and the experiments designed by W.C. The TAB-seq and oxBS-seq library construction and genome analyses were performed by Q.M. Z.X., H.L., and Z.X., and Y.Z. were involved in tissue sampling. Y.B. performed the mass spectrometry experiments.

